# NUDT6, the Antisense Protein of FGF2 Gene, Plays a Depressogenic Role by Promoting Inflammation and Suppressing Neurogenesis without Altering FGF2 Signaling

**DOI:** 10.1101/2022.09.05.506638

**Authors:** Burak Uzay, Fatma Özlem Hökelekli, Murat Yılmaz, Emre Cem Esen, Koray Başar, Aslıhan Bahadır-Varol, Yavuz Ayhan, Turgay Dalkara, Emine Eren-Koçak

**Affiliations:** Institute of Neurological Sciences and Psychiatry, Hacettepe University, Ankara, Turkey; Brain Institute, Department of Pharmacology, Vanderbilt University, Nashville, 37240-7933,TN, USA; Department of Psychiatry, University of Texas Southwestern, Dallas, TX, USA; Department of Biology, Faculty of Science, Hacettepe University, Ankara, Turkey; Department of Psychiatry, Faculty of Medicine, Hacettepe University, Ankara, Turkey

**Author notes:** These authors contributed equally to this work (Fatma Özlem Hökelekli, Murat Yılmaz). Corresponding author, Corresponding author: Emine Eren-Kocak, Hacettepe Universitesi, Norolojik Bilimler ve Psikiyatri Enstitusu, Tip Fakultesi Binasi, Asma kat, Sihhiye, 06100 / Ankara/ Turkey.

**Keywords:** NUDT6, FGF2, FGF-AS, natural antisense transcripts, depression, S100A9, NF-κB, neurogenesis

## Abstract

Fibroblast growth factor-2 (FGF2) is involved in the regulation of affective behavior and shows antidepressant effects through Akt and ERK1/2 pathways. NUDT6 is a protein encoded from FGF2 gene’s antisense strand and its role in the regulation of affective behavior is unclear. Here, we show that increasing NUDT6 expression in the hippocampus results in depression-like behavior in rats without changing FGF2 levels or activating its downstream effectors, Akt and ERK1/2. Instead, NUDT6 acts by inducing inflammatory signaling, specifically by increasing S100A9 levels, activating NF-κB and rising microglia number along with a reduction in neurogenesis. Conversely, inhibition of hippocampal NUDT6 expression by shRNA results in antidepressant effects and increases neurogenesis without altering FGF2 levels. Together these findings suggest that NUDT6 may play a role in major depression by inducing a proinflammatory state and serve as a novel therapeutic target for antidepressant development. This is the first report of an antisense protein acting through a different mechanism of action than regulation of its sense protein. The opposite effects of NUDT6 and FGF2 on depression-like behavior may serve as a mechanism to fine-tune affective behavior. Our findings open up new venues for studying the differential regulation and functional interactions of sense and antisense proteins in neural function and behavior as well as in neuropsychiatric disorders.

## Introduction

Major depressive disorder (MDD) is a debilitating psychiatric disorder affecting 4.7% of the world population (Ferrari et al., 2013). Despite several treatment options, treatment resistance affecting approximately one third of patients is a major concern (Nemeroff, 2007; Trivedi et al., 2006). Current treatment strategies for major depression lack specificity in targeting particular pathophysiological mechanisms. Therefore, better understanding of the pathophysiological mechanisms underlying MDD is necessary to develop new antidepressant interventions.

One of the leading hypotheses of major depression is the “neurotrophic hypothesis”. It stems from observations of reduced growth factor levels in several brain regions of chronically stressed animals and human depressed subjects, which are reversed by antidepressant treatments (reviewed in (Duman and Monteggia, 2006; Turner et al., 2012)). In addition, increasing brain derived neurotrophic factor (BDNF) and fibroblast growth factor-2 (FGF2) levels in brain has antidepressant and anxiolytic effects in rodent models (Eren-Koçak et al., 2011; Hoshaw et al., 2008; Perez et al., 2009; Turner et al., 2008). Both FGF2 and BDNF have natural antisense transcripts (NAT), RNA molecules that are transcribed from the opposite strand of protein-coding genes, and are suggested to regulate the transcription of their sense strand (Modarresi et al., 2012). Supporting this, BDNF-antisense (BDNF-AS) is reported to be a discordant regulator of BDNF protein expression i.e. when BDNF-AS levels increase, BDNF protein expression decreases (Xu et al., 2022). A similar role was suggested for FGF-AS in the regulation of FGF2 expression based on initial *in vitro* studies (Baguma-Nibasheka et al., 2012, 2007; Li and Murphy, 2000; MacFarlane et al., 2010), however several other *in vitro* studies failed to detect an interaction between FGF-AS and FGF2 expression levels (Asa et al., 2001; Baguma-Nibasheka et al., 2012, 2005; MacFarlane et al., 2010). FGF-AS is translated into a 35 kDa protein, nudix hydrolase 6 (NUDT6), the function of which is unknown (Asa et al., 2001). Intriguingly, hippocampal NUDT6 expression was altered in response to acute stress in rats, suggesting a role for NUDT6 in the regulation of affective behavior (Frank et al., 2007; Xu et al., 2017). Although this observation points to an additional layer of complexity in the regulation of the affective behavior by neurotrophic factors, the place of NUDT6 in MDD pathophysiology has not yet been elucidated. Understanding the potential role of NUDT6 in affective behavior may therefore provide a deeper insight to mechanisms of MDD and disclose novel antidepressant therapeutic targets (Elsayed et al., 2012; Eren-Koçak et al., 2011; Perez et al., 2009).

Here, we show that overexpression of hippocampal NUDT6 increases anxiety- and depression-like behaviors in rats. These effects were independent of FGF2 signaling, as we did not find any changes in hippocampal FGF2 expression or its downstream signaling mediated by Akt and ERK1/2. Rather, hippocampal NUDT6 overexpression increased 1) the expression of proinflammatory S100A9, an alarmin / damage associated molecular pattern (DAMP), 2) nuclear translocation (thus activation) of NF-κB-p65, a proinflammatory transcription factor and 3) number of microglia, all conforming to an inflammatory state. Moreover, NUDT6 overexpression reduced neurogenesis, possibly as a consequence of the proinflammatory changes induced (Koo et al., 2010). Confirming the role of NUDT6 in depression pathophysiology, we found that its knockdown in the hippocampus not only had antidepressant and anxiolytic effects, but also stimulated neurogenesis. Overall, our results suggest that NUDT6 can promote inflammatory signaling in the hippocampus, which may take part in decreased neurogenesis and depression-like behavior observed on its overexpression and, reversed by its knockdown. Our study discloses a novel mechanism in the pathophysiology of MDD and, for the first time, underscores the functional effects of antisense proteins in neuropsychiatric diseases. The fact that NUDT6 regulates affective behavior through a different mechanism than its sense transcript, FGF2, whose antidepressant effects is well known, may open up new venues for studying the functional interactions of sense and antisense proteins not only in MDD but also in other CNS disorders.

## Results

### 1. Hippocampal overexpression of NUDT6 induces depression- and anxiety-like behaviors

To determine whether hippocampal NUDT6 is involved in the regulation of affective behavior, first we overexpressed NUDT6 in the hippocampus by two means: 1. Repeated daily intracerebroventricular (ICV) injections of an NUDT6-expressing plasmid (NUDT6-EP) for 14 days to avoid hippocampal tissue injury and growth factor response that can result from site specific microinjection (Turner et al., 2016), 2. Bilateral intrahippocampal microinjection of NUDT6-expressing adeno-associated virus (NUDT6-AAV2) to ensure hippocampus-specific overexpression (Figure 1A). At the end of the ICV injections and one month following the microinjection of NUDT6-AAV2, we performed behavioral tests to assess depression-like behavior by sucrose preference test (SPT) and forced swim test (FST), and anxiety-like behavior by elevated plus maze test (EPM) (Figure 1A). Both repeated ICV injections of NUDT6-EP and bilateral intrahippocampal injections of NUDT6-AAV2 increased immobility time and decreased swimming time in FST, in accordance with an increase in depression-like behavior (p= 0.067 and 0.050 for ICV injections, and 0.005 and 0.003 for AAV2 injections, respectively) (Figure 1B and 1C). There was, however, no change in SPT in the NUDT6-EP or NUDT6-AAV2 groups compared to their corresponding controls treated with vehicle or Blank-AAV (p=0.501 and 0.617, respectively) (Figure S1A and S1B).

**Figure 1.**
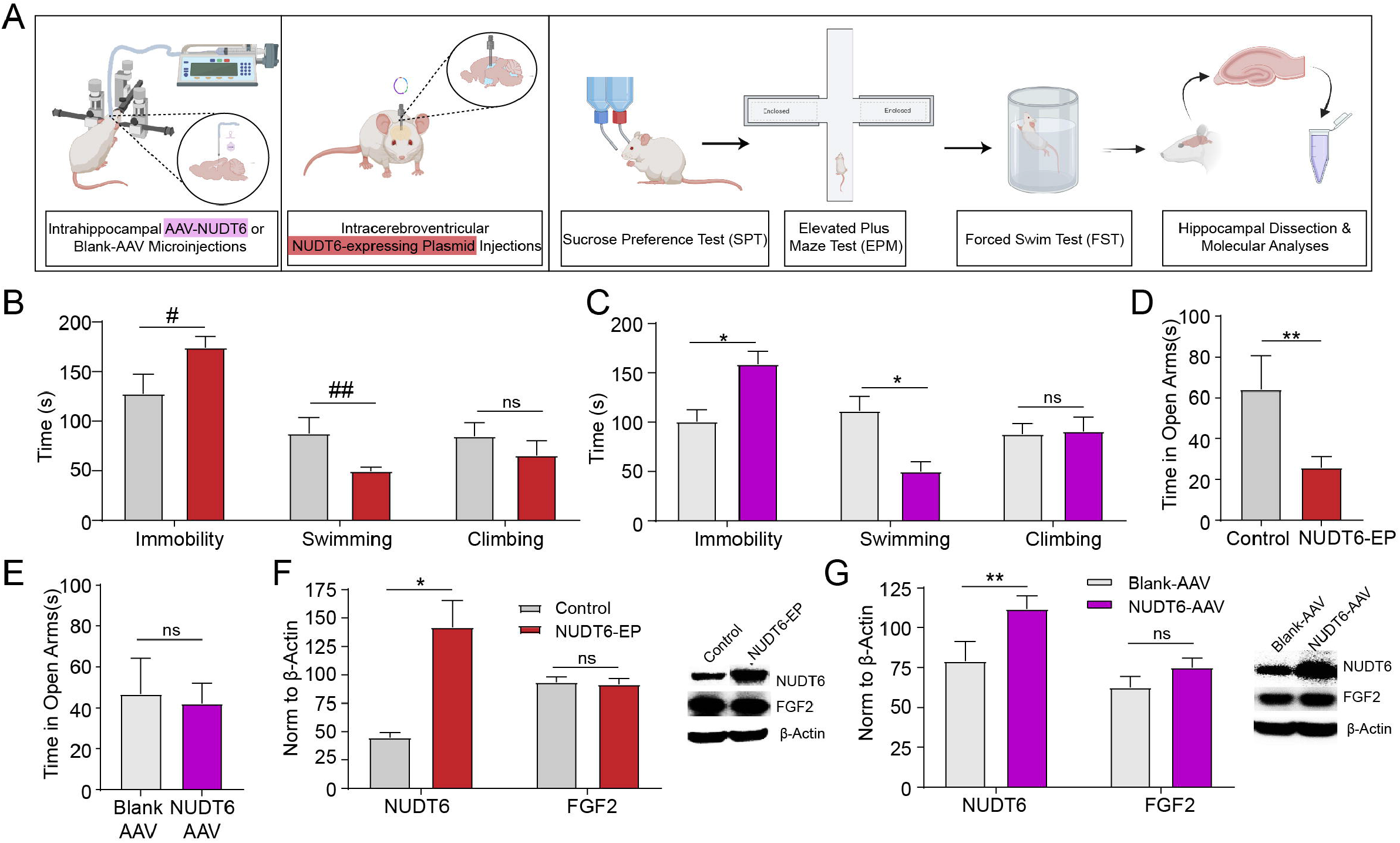
NUDT6 overexpression in the hippocampus increases depression- and anxiety-like behaviors. **A.** Graphical depiction of the NUDT6-EP and NUDT6-AAV experimental setup. **B.** Repeated icv injections of NUDT6-EP increased time spent immobile and decreased time spent swimming in forced swim test, when compared to controls (n=7-11) **C**. Intrahippocampal injection of NUDT6-AAV increased time spent immobile and decreased time spent swimming in forced swim test, when compared to Blank-AAV injected controls (n=13/group). **D.** Repeated icv injections of NUDT6-EP decreased time spent in open arms in elevated plus maze (n=7-11/group). **E.** Intrahippocampal injection of NUDT6-AAV did not alter time spent in open arms in elevated plus maze (n=13/group). **F.** Repeated icv injections of NUDT6-EP increased hippocampal NUDT6 protein levels. Hippocampal FGF2 protein levels, on the other hand, remained unchanged. Representative Western bands for NUDT6 and FGF2 of each group are shown; ß-actin is used as loading control (n=5-6/group). **G.** Intrahippocampal NUDT6-AAV injection increased NUDT6 mRNA expression and protein levels. FGF2 mRNA expression or protein levels in the hippocampus were not altered as detected by RT-PCR and Western blotting (n=7-8/group). Representative Western bands for NUDT6 and FGF2 of each group are shown; ß-actin is used as loading control. FGF2: fibroblast growth factor 2, NUDT6-AAV: NUDT6 expressing adeno-associated virus, NUDT6-EP: NUDT6-expressing plasmid, RT-PCR: real-time polymerase chain reaction. Significance levels were stated as follows: #p=0.067, ##p=0.05, *p < 0.05, **p < 0.01, ***p < 0.001, and ****p < 0.0001. ns denotes non-significance.

NUDT6-EP group spent less time in open arms in EPM than the vehicle-injected control group, indicating an increase in anxiety-like behavior (p=0.032) (Figure 1D). Time spent in open arms in EPM was similar in the NUDT6-AAV2 and Blank-AAV2 injected groups (p=0.89)(Figure 1E). The total distance moved in EPM was also similar between experimental and control groups in both sets of experiments (p=0.274 and 0.451, respectively) (Figure S1C and S1D).

### 2. Altering NUDT6 expression neither changes FGF2 levels nor interferes with FGF2 signaling pathways in the hippocampus

We investigated whether the effects of NUDT6 on depression- and anxiety-like behavior were mediated by a change in the expression of its sense transcript, FGF2. After confirming the increase in NUDT6 protein levels in the hippocampus in both groups by immunoblotting (p=0.009 and p=0.04 in ICV NUDT6-EP, intrahippocampal NUDT6-AAV2 injection groups, respectively), we examined the corresponding hippocampal FGF2 protein levels (Figures 1F and 1G). On contrary to *in vitro* studies reporting transcriptional regulation of FGF2 by FGF-AS (Baguma-Nibasheka et al., 2012, 2007; MacFarlane et al., 2010), we found no change in FGF2 protein levels by increasing NUDT6 expression (p=0.80 and p=0.20 in ICV NUDT6-EP and intrahippocampal NUDT6-AAV2 injection groups, respectively) (Figures 1F and 1G). FGF2 mRNA expression was also not changed after intrahippocampal NUDT6-AAV injections (p=0.59), ruling out a possible regulatory effect of NUDT6 on FGF2 transcription (Figure S1F).

The above findings strongly suggest that the depressogenic effect of hippocampal NUDT6 was not mediated by a change in FGF2 levels. To rule out possible interactions at downstream mediator levels, we studied the phosphorylation of ERK1/2 and Akt in NUDT6-AAV2/Blank AAV2 microinjected hippocampi, molecules mediating the antidepressant actions of FGF2 (Tang et al., 2017; Wang et al., 2018) (Figure 2A). We did not find any difference in the phosphorylation of either Akt or ERK1/2 between the Blank-AAV2 control and NUDT6-AAV2 groups (p>0.99 and p=0.80 respectively) (Figure 2B and 2 C).

**Figure 2.**
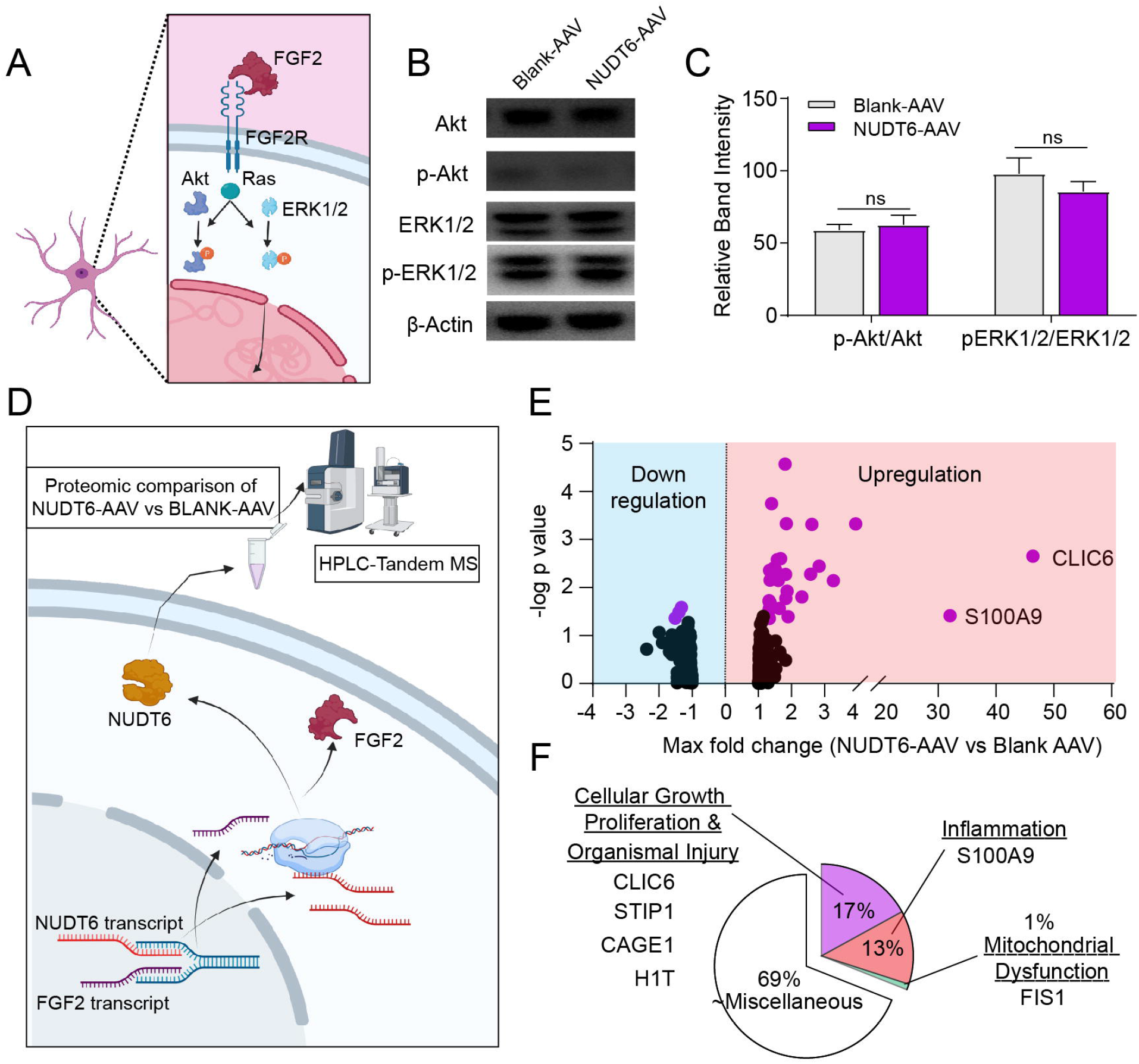
NUDT6 overexpression in the hippocampus did not cause any change in either FGF2 levels or phosphorylation of its downstream signaling pathways Akt and ERK1/2. **A.** Graphical depiction showing FGF2 signaling pathways. **B.** Representative Western blot bands for Akt, p-Akt, ERK1/2, p-ERK1/2 of each group are shown; β-actin is used as a loading control (n=7/group) **C.** NUDT6 overexpression did not change phosphorylation of Akt or ERK1/2 proteins shown by relative band densities (n=7/group) (ns denotes non-significance). **D.** Graphical depiction of the HPLC-Tandem MS proteomics experimental setup. **E.** Proteomic analysis revealed an upregulation of 31 and a downregulation of 3 proteins in the NUDT6-AAV group (n/7/group). **F.** Thirteen percent (13%) of the upregulated proteins were related to inflammatory signaling whereas 1% were related to mitochondrial dysfunction and 17% were related to cellular growth & proliferation and organismal injury. The remaining 69% is related to miscellaneous pathways. FGF2: fibroblast growth factor 2, NUDT6-AAV: NUDT6 expressing adeno-associated virus, HPLC-Tandem MS: High performance Liquid Chromatography Mass Spectrometry

### 3. Increasing NUDT6 expression induces inflammatory signaling

To have an insight into the molecular mechanisms of action of NUDT6, we performed proteomic analysis of hippocampal samples from NUDT6-AAV2 and Blank-AAV2 groups (Figure 2D), which revealed a significant upregulation of 31 proteins and downregulation of 3 proteins in the NUDT6-AAV2 group compared to controls (p<0.05) (Figure 2E) (Supplemental Table 1). When these 34 proteins were subjected to in silico pathway analysis using online Panther and String softwares, we found that they are involved in neuroinflammation, cellular growth and proliferation, organismal injury and abnormalities and mitochondrial dysfunction (Figure 2F). FGF2 was not among the 34 proteins that were changed by hippocampal NUDT6 overexpression in line with the above immunoblotting and RT-PCR findings. However, there was a 32-fold increase in the expression of S100 calcium binding protein A9 (S100A9), a proinflammatory molecule mainly expressed by myeloid cells (i.e. microglia in brain), supporting the role of inflammatory signaling in the development of NUDT6-induced depression-like behavior (Iwata et al., 2016; Koo et al., 2010) (Figure 2E).

Western blotting and RT-PCR of hippocampal samples confirmed the increase in S100A9 mRNA and protein levels in the NUDT6–AAV group compared to Blank-AAV control group (p=0.04 and p=0.01, respectively) (Figure 3A-C). We double-immunostained the hippocampal sections from NUDT6-AAV and Blank-AAV groups with either anti-S100A9 & anti-NeuN antibodies or with anti-S100A9 & anti-Iba1 antibodies to label neuronal or microglial S100A9, respectively. We found that S100A9 was colocalized to both neurons and microglia and its immunopositivity in both dentate granular and hilar neurons was significantly increased in NUDT6-AAV group relative to the Blank-AAV control group (p<0.0001) (Figure 3D-E and Figure S2A). To test whether S100A9 was increased in the same cells in which NUDT6 was overexpressed, we double-immunostained hippocampal sections with NUDT6 &S100A9. There was a significant increase in S100A9 in NUDT6-positive cells in the NUDT6-AAV2 group compared to Blank-AAV2 group, suggesting that the increase in S100A9 took place downstream to NUDT6 in the same cell (Figure 3F-G).

**Figure 3.**
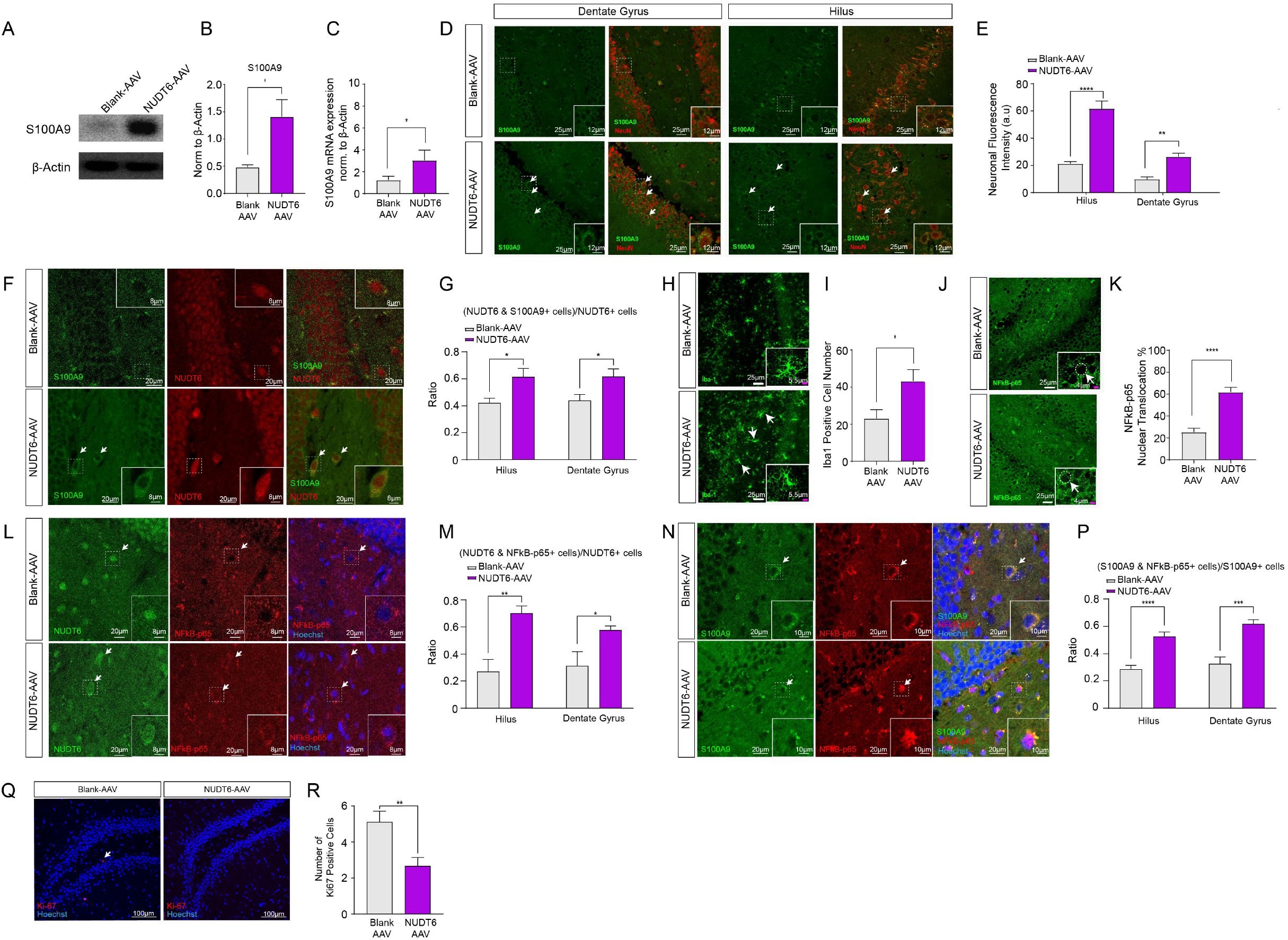
NUDT6 overexpression in the hippocampus promotes inflammatory signaling. **A-B.** NUDT6 overexpression increased S100A9 protein expression. Representative Western bands for S100A9 of each group are shown; ß-actin is used as loading control (n =7/group). **C.** NUDT6 overexpression increased S100A9 mRNA expression (n =7-8/group). **D.** Colocalization of S100A9 with a neuronal cell marker, NeuN showed that S100A9 levels were increased in neurons. Neurons were labeled with NeuN (red) and S100A9 was labeled with green. Scale bar: 25 μm. **E.** Total florescence of S100A9 in NeuN positive cells was increased in both the dentate gyrus and the hilus in the NUDT6-AAV group, when compared to the control group (n =3/group). **F-G.** Colocalization of S100A9 and NUDT6 showed that S100A9 significantly increased in NUDT6-positive cells in NUDT6-AAV group compared to the control group. (n =3/group). Scale bar: 20 μm. **H-I.** Number of Iba1 labeled cells in the dentate gyrus and hilus was higher in the NUDT6-AAV group than the control group (n =3/group). Scale bar: 25 μm and 5.5 μm for the inset. **J-K.** Nuclear translocation of NF-κB was increased in the NUDT6-AAV group in the dentate gyrus (n =3/group). Figure insets show representative images of nuclear labeling of NF-κB (indicating translocation) in dentate gyrus. Scale bar: 25 μm and 4 μm for the inset. **L-M.** Colocalization of NF-κB and NUDT6 showed that nuclear translocation of NF-κB significantly increased in NUDT6-positive cells in NUDT6-AAV group compared to the control group (n =3/group). Scale bar: 20 μm. **N-P.** Colocalization of S100A9 and NF-κB showed that NF-κB activation was significantly higher in S100A9-positive cells in NUDT6-AAV group than the control group (n =3/group). Scale bar: 20 μm. **Q-R.** Hippocampal Ki-67 immunofluorescent staining showed that neurogenesis was significantly lower in NUDT6-AAV group than the control group (n =3/group). Scale bar: 100 μm. Significance levels were stated as follows: *p < 0.05, **p < 0.01, ***p < 0.001, and ****p < 0.0001. ns denotes non-significance.

As S100A8/A9 reportedly results in microglial cell migration *in vitro* (23), we assessed whether NUDT6 overexpression altered the number of Iba-1 immunoreactive cells (microglia) in hippocampus (Ma et al., 2017). The number of Iba-1 labeled microglia was increased by %87 in hilar region of the hippocampus in NUDT6-AAV2 group compared to Blank-AAV2 group, which indicates microglial recruitment in response to NUDT6 overexpression and further supports the involvement of inflammatory signaling in the NUDT6’s depressogenic actions (p=0.01) (Figure 3H-I).

Previous studies report that S100A9 is associated with the activation of proinflammatory transcription factor NF-κB (Németh et al., 2009; Vogl et al., 2007). When NF-κB binding protein IκB is phosphorylated, the p65 subunit of NF-κB is freed and translocated to the nucleus, where it initiates transcription of several inflammatory molecules, including cytokines and chemokines (Shih et al., 2015). Thus, nuclear translocation of NF-κB-p65 is widely used as an indicator of increased neuroinflammatory signaling (Eren-Koçak and Dalkara, 2021; Liu et al., 2017). We investigated whether NUDT6 overexpression induced activation of NF-κB-p65 in our samples by quantifying the ratio of NF-κB-p65 positive nuclei-to-total nuclei in immunostained hippocampal sections from NUDT6-AAV2 and blank-AAV2 groups. We found a 2.4-fold increase in NF-κB nuclear translocation in the dentate gyrus of NUDT6-AAV group compared to Blank-AAV injected controls (p<0.0001), indicating activation of NF-κB by hippocampal NUDT6 overexpression (Figure 3J-K). Double labeling of NF-κB-p65 with NUDT6 or S100A9 revealed an increased NF-κB-p65 nuclear translocation in both NUDT6- and S100A9-positive cells in the NUDT6-AAV2 group compared to Blank-AAV2 group, respectively (p=0.0021 for hilus and p=0.031 for dentate gyrus, p<0.0001 for hilus and p=0.0003 for dentate gyrus, respectively) (Figure 3L-P).

### 4. Overexpression of NUDT6 reduces hippocampal neurogenesis

Due to the putative role of neurogenesis in the regulation of affective behavior (Dranovsky and Hen, 2006; Hill et al., 2015; Santarelli et al., 2003) and involvement of NF-κB in the regulation of in hippocampal neurogenesis (Koo et al., 2010), we investigated the effect of NUDT6 overexpression on hippocampal neurogenesis by Ki-67 immunostaining, a marker for cell proliferation (Figure 3Q). The total number of Ki-67-positive cells was significantly decreased in NUDT6-AAV2 group compared to Blank-AAV2 group (p=0.04) (Figure 3R). These findings suggest that inhibition of neurogenesis may be a potential contributor to NUDT6’s mechanism of action in mediating depression-like behavior.

### 5. Hippocampal NUDT6 knockdown reduces depression- and anxiety-like behavior and increases hippocampal neurogenesis

To determine if NUDT6 could be a potential target for antidepressant interventions, we knocked down the expression of NUDT6 by bilateral intrahippocampal microinjections of lentivirus expressing NUDT6-shRNA (Figure 4A). One month after the injections, we performed the behavioral tests FST and SPT to assess depression-like behavior and EPM test to assess anxiety-like behavior. Following behavioral experiments, the rats’ hippocampi were dissected and analyzed for NUDT6 and FGF2 protein levels, in addition to Ki-67 immunostaining to evaluate neurogenesis (Figure 4A). There was a significant decrease in NUDT6 protein levels in the NUDT6-shRNA group compared to the non-targeting-shRNA group confirming a successful knockdown (Figure 4B). We did not observe any difference in FGF2 levels between groups (Figure 4B). Hippocampal NUDT6 knockdown resulted in an increase in sucrose preference in the SPT when compared to the non-targeting-shRNA control group (p=0.026) (Figure 4C). On the other hand, we did not observe any differences in durations spent immobile, swimming and climbing in FST between groups (p=0.99, 0.83 and 0.72, respectively) (Figure 4D).

**Figure 4.**
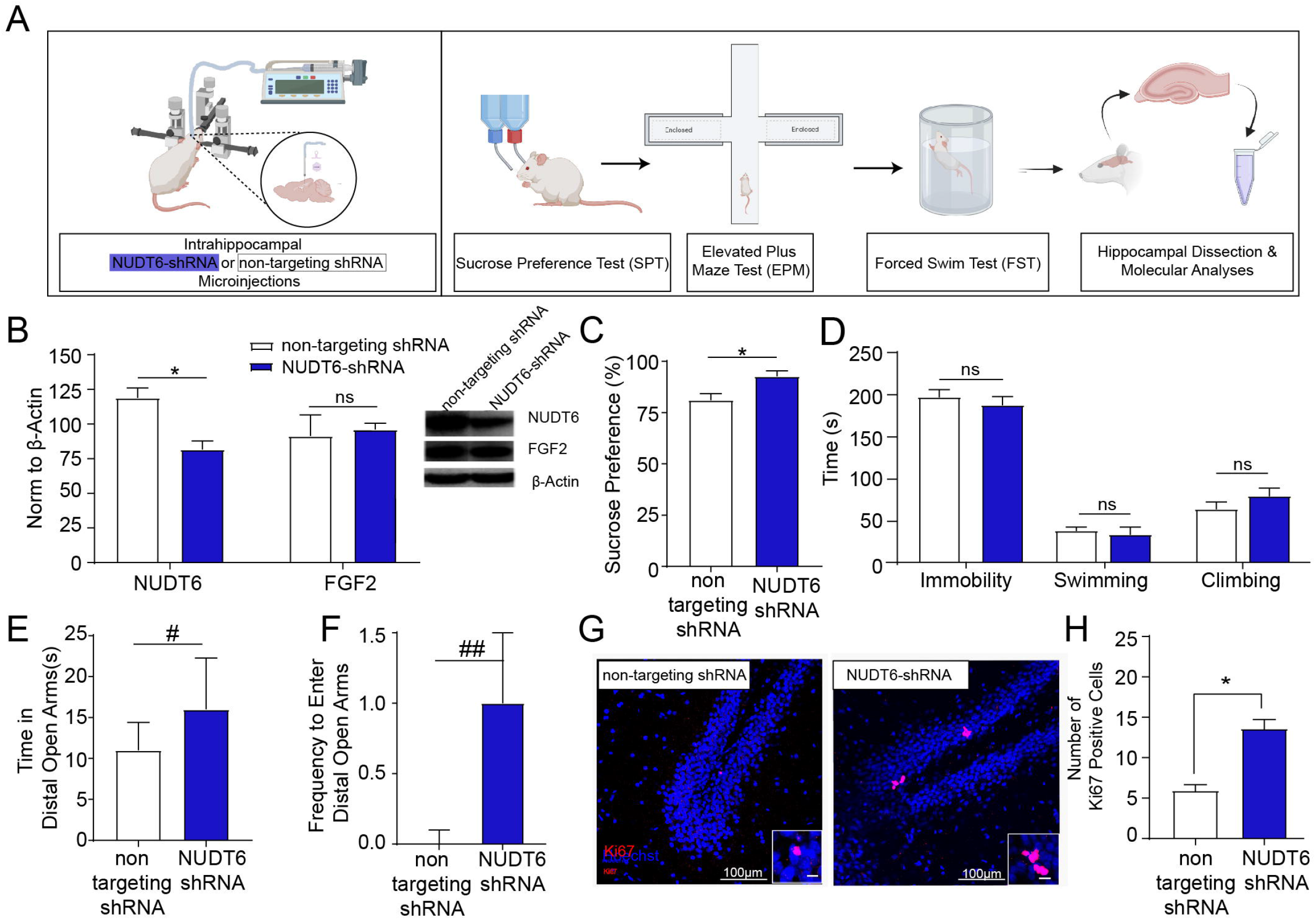
Hippocampal NUDT6 knockdown reduces depression- and anxiety-like behavior and increases hippocampal neurogenesis **A.** Schematic representation of experimental setup of hippocampal NUDT6 knockdown via NUDT6-shRNA **B.** Intrahippocampal NUDT6-shRNA injection decreased NUDT6 protein expression relative to the non-targeting shRNA injected group. Representative Western bands for NUDT6 and FGF2 of each group are shown; β-actin is used as loading control (n=5-6/group). **C.** NUDT6 knockdown by intrahippocampal NUDT6-shRNA injection resulted in an increase in sucrose preference (n=6-7/group). **D.** Knockdown of NUDT6 in the hippocampus did not change time spent immobile, climbing or swimming in the forced swim test when compared to non-targeting-shRNA control group. **E-F.** NUDT6-shRNA group spent more time and entered more frequently into the distal open arms of elevated plus maze (n=11-13/group). **G-H.** Hippocampal Ki-67 immunofluorescent staining showed that neurogenesis was significantly increased in NUDT6-shRNA group compared to the non-targeting shRNA group (n =3/group). Scale bar: 100 μm. NUDT6-shRNA: NUDT6 short hairpin RNA. Significance levels were stated as follows: #p=0.10, ##p=0.14, *p < 0.05, **p < 0.01, ***p < 0.001, and ****p < 0.0001. ns denotes nonsignificance.

Experimental reduction of hippocampal NUDT6 protein levels increased total time spent in distal half of the open arms as well as the frequency of entrance into the distal open arms in EPM, indicating an overall anxiolytic effect (p=0.10 and 0.14, respectively) (Figure 4E-F). Total distance traveled in the EPM was similar between groups (p=0.86), suggesting that the difference reported above was not due to a change in locomotor activity (Figure S1E).

Finally, we evaluated the effects of hippocampal NUDT6 knockdown on neurogenesis by Ki-67 immunostaining, and found a significant increase in Ki-67-positive cell number in NUDT6-shRNA group (Figure 4G, 4H). In summary knockdown of NUDT6 in the hippocampus had antidepressant and anxiolytic effects and was associated with an increase in neurogenesis, identifying NUDT6 as a potential target in developing novel antidepressants.

## Discussion

In this study, we showed that increasing hippocampal NUDT6 levels results in depression-like behaviors by inducing inflammatory signaling and suppressing hippocampal neurogenesis. HPLC-MS proteomics analysis showed that hippocampal NUDT6 overexpression led to increased expression of S100A9, a DAMP/alarmin, which was confirmed by immunoblotting and qRT-PCR. Immunostaining for S100A9 increased by approximately 3 times in hilar and dentate gyrus neurons. Importantly, S100A9 positive cells in hilus and dentate gyrus exhibited a 2-3 fold rise in nuclear translocation of NF-κB, the pro-inflammatory transcription factor. In parallel with S100A9 rise and NF-κB activation, microglial cell number was found to be increased. Microglia are likely to be recruited to the hilar region in response to chemokines released because of the proinflammatory state induced by NUDT6 overexpression. These findings suggest that hippocampal NUDT6 overexpression leads to depression-like behavior possibly by promoting a proinflammatory state, which is highly regarded to play an important role in depression pathophysiology based on several lines of evidence in the literature (Bierhaus et al., 2003; Gárate et al., 2013; Gong et al., 2018; Koo et al., 2010; Wachholz et al., 2016; Wohleb et al., 2018). To our knowledge, this is the first direct evidence showing that overexpression of a non-inflammatory protein in the hippocampus can induce neuroinflammation and depression-like behavior in naïve mice, strongly supporting the neuroinflammatory theory of depression.

We did not find any changes in FGF2 mRNA or protein levels after NUDT6 overexpression in the hippocampus, on contrary to previous *in vitro* evidence from cancer cells, suggesting negative regulation of FGF2 expression by the FGF-AS transcript (Baguma-Nibasheka et al., 2012, 2007; Knee et al., 1997; Li and Murphy, 2000; MacFarlane et al., 2010). This discrepancy may have resulted from completely different gene regulatory mechanisms in highly mitotic cancer cells and post-mitotic brain cells. Supporting this suggestion, Asa et al (2001) reported that FGF-AS overexpression did not alter FGF2 expression in a pituitary derived cell line (Asa et al., 2001). In addition, we found that NUDT6 overexpression did not change phosphorylation of ERK1/2 and Akt kinases, the downstream signaling mediators of FGF2 that mediate its antidepressant effects (Tang et al., 2017; Wang et al., 2018). Thus, we conclude that the effects of the NUDT6 protein in the hippocampus on affective behavior are not mediated by modulation of FGF2 or its downstream signaling pathways, but NUDT6 has functions of its own.

In line with our findings, involvement of S100A9 in stress response and depression-like behavior has been reported by a recent study, which showed that chronic unpredictable mild stress (CUMS), a well-established depression model increased hippocampal S100A9 levels (Gong et al., 2018). In this study, ICV administration of recombinant S100A9 increased depression-like behavior in mice, whereas intraperitoneal injection of S100A9 inhibitor during CUMS period blocked depressogenic effects of CUMS in mice (Gong et al., 2018). The S100 protein family is comprised of several calcium-binding proteins, which form homo-/ hetero-dimers that function by crosslinking target proteins (Donato, 2001). This way, they take part in the regulation of various cellular functions, including inflammation, cell proliferation and differentiation, cytoskeletal dynamics, and transcription (Donato, 2003, 2001). In addition to their intracellular functions, some S100 proteins, including S100A9, are secreted by unknown mechanisms, which then bind to pattern recognition receptors on the cell membrane resulting in NF-κB activation in the target cell (Chen et al., 2015; Donato, 2003; Ma et al., 2017). S100A9 can also induce cytokine and chemokine secretion from microglial cell cultures as well as from peripheral blood monocytic cells, which can activate NF-κB in neighboring cells (Simard et al., 2013; Wu et al., 2018). Consistent with these reports, we found an increase in nuclear translocation of NF-κB along with increased levels of S100A9 in the NUDT6 overexpression group. We further showed that S100A9 immunopositivity was increased in NUDT6-positive cells, and NF-κB activation was increased in both NUDT6-positive and S100A9-positive cells. These colocalizations suggest that NUDT6 overexpression induces neuronal S100A9 and activates NF-κB. The consequent NF-κB activation could result in increased transcription and release of cytokines and chemokines, augmenting the proinflammatory state. Supporting these suggestions, stress has been shown to promote NF-κB activation in both hippocampus and prefrontal cortex, and ICV administration of an NF-κB inhibitor during 4 weeks of CUMS blocked the effects of CUMS on anxiety- and depression-like behavior in mice (Gárate et al., 2013; Koo et al., 2010).

We observed an increase in the number of microglia in the hippocampus after NUDT6 overexpression. This finding is consistent with previous reports of increased number of microglia in animal models of depression (Wachholz et al., 2016; Wohleb et al., 2018). In addition, S100A9 has been reported to promote microglial cell migration in microglial cell cultures (Wu et al., 2018). The recruitment of microglial cells by S100A9 or chemokines transcribed by activated NF-κB may explain the observed increase in the number of microglia after NUDT6 overexpression.

Overexpression of NUDT6 resulted in reduced hippocampal neurogenesis, which may be one of the pathways mediating the depressogenic effect of NUDT6. This finding conforms to a vast body of literature reporting the involvement of adult hippocampal neurogenesis in depression pathophysiology (reviewed in (Chooniedass-Kothari et al., 2004)). Briefly, hippocampal neurogenesis was reportedly suppressed by both acute and chronic stress paradigms modeling depression in rodents (Gascoigne et al., 2012; Stergachis et al., 2013) (reviewed in (Haapakoski et al., 2015)), which was reversed by antidepressant treatments (Santarelli et al., 2003; Taupin, 2006). Furthermore, the effects of acute and chronic stress on neurogenesis have been reported to involve NF-κB signaling. Stress has been shown to activate NF-κB in the adult hippocampus, while inhibition of NF-κB signaling blocked the effects of acute or chronic stress on neurogenesis, as noted above (Koo et al., 2010). Our findings of decreased neurogenesis together with activation of NF-κB in NUDT6 overexpression group suggest that NUDT6 may suppress neurogenesis by activating inflammatory pathways. In summary, NUDT6 overexpression induces proinflammatory activity in the hippocampus, possibly by promoting the synthesis of chemokines and cytokines, which then inhibit neurogenesis and recruit microglia.

Although the effects of increasing NUDT6 expression by plasmid (NUDT6-EP) or virus (NUDT6-AAV2) transfection on depression-like behavior were similar that was not the case for anxiety-like behavior. Whereas icv administration of NUDT6-EP was clearly anxiogenic, NUDT6-AAV2 hippocampal injection had no observable effects of on anxiety-like behaviors. This difference can be attributed to higher NUDT6 expression achieved by NUDT6-EP (142 vs. 33%). NUDT6 knockdown by 69% also affected anxiety-like behavior, supporting this view. Intriguingly, depending on whether NUDT6 expression was promoted or suppressed, depression-related behavioral tests exhibited partly discordant outcomes rather than changes in expected (i.e., opposite) directions. One possible explanation is engagement of a different molecular mechanism by NUDT6 knockdown from suppression of inflammatory signaling. Thus, NUDT6 overexpression and NUDT6 knockdown may affect different aspects of depression-like behavior by affecting different molecular pathways. It is also possible that the amplifying nature of the inflammatory cascade triggered by NUDT6 overexpression leads to more pronounced behavioral changes than NUDT6 suppression. Indeed, we showed that several players of the inflammatory cascade were activated by NUDT6 overexpression. Another plausible explanation may be the use of naïve animals in the NUDT6 knockdown experiments. As naïve rats do not display increase in depression-like behaviors, the antidepressant effect of knocking down NUDT6 could not be unmasked in these rats. This possibility should be addressed by future studies designed to study the rescue effects of NUDT6 knockdown in a depression model.

We found that hippocampal NUDT6 knockdown had antidepressant and anxiolytic effects, identifying NUDT6 as a novel antidepressant target. These effects of NUDT6 knockdown were accompanied by an increase in hippocampal neurogenesis. As neurogenesis has been reported to be necessary for behavioral effects of the antidepressants (Santarelli et al., 2003), our findings indicate that prevention of suppressive effects of NUDT6 on neurogenesis may be involved in antidepressant and anxiolytic effects of NUDT6 knockdown. These effects, similar to those of NUDT6 overexpression, were not associated with changes in hippocampal FGF2 levels.

In conclusion, our results identify NUDT6 as a potential mediator of depression-like behavior and a novel target for the antidepressant drug development. NUDT6’s depressogenic effect was not associated with a change in FGF2 signaling, rather it was associated with induction of neuroinflammatory signaling, which involves S100A9 expression, NF-κB activation and microglial recruitment. These findings are consistent with previous reports that suggest a role for inflammatory signaling in the development of depression (Gong et al., 2018; Haapakoski et al., 2015; Koo et al., 2010; Wachholz et al., 2016). Our findings add NUDT6 and S100A9 among the pro-inflammatory players in depression pathophysiology and reveal novel targets for MDD treatment. Furthermore, this is the first *in vivo* study, to our knowledge, to disclose a functional role for an antisense protein in the CNS, independent of the levels of its sense protein. These findings suggest that transcription of the same gene in the sense and antisense direction may enable the organism to fine-tune behavioral responses as exemplified here by NUDT6 and FGF2, both transcribed from FGF2 gene but have opposite actions on depression-like behavior. This encouraging finding may lead to new discoveries uncovering other protein-coding NATs that may have biological functions independent of the regulation of their sense transcript.

As for the limitations, we performed the experiments in only male rats, therefore our findings cannot be generalized to females. There is evidence in the literature that microglia and neurons respond differentially to physical/psychological stressors in male animals than in females (Cao et al., 2021; Wohleb et al., 2018). Therefore, possible involvement of NUDT6 in the development of depression-like behavior in females needs to be addressed in future studies.

## Methods and Materials

### ANIMALS

Male Sprague Dawley rats, weighing 300 to 400 grams were used in this study. All animals were maintained on a 12-hour light/dark cycle with food and water available ad libitum. Animals were habituated to the housing conditions for 1 week before any manipulations. They were weighed twice weekly and monitored closely for any deterioration in their health. All the experimental procedures were approved by the Animal Experimentations Local Ethics Board of the Hacettepe University.

### PLASMIDS

pEGFP-N1 plasmid expressing full-length rat FGF-AS gene (a generous gift from Dr. Paul Murphy, Dalhousie University, Canada) was purified as endotoxin-free plasmids (Qiagen, endo-free plasmid Maxi-kit). To facilitate the in vivo delivery of the plasmids, a transfection reagent was used (in vivo-JetPEI-TM, Polyplus transfection France) according to the manufacturer’s protocol. Briefly, 3 μg of plasmid was mixed with in vivo JetPEI with an N/P ratio of 6 in a total volume of 7 μl for each rat. The plasmid-in vivo JetPEI mix were prepared daily, 30 minutes before the microinjections. For the controls, we used in vivo JetPEI mix prepared without the plasmid.

### VIRUS

Adeno-associated virus (AAV) serotype 2 was used to overexpress full-length rat NUDT6 in the hippocampus (pAAV-G-CMV-2A-GFP)(Abmgood, aavp6966983). A blank AAV (pAAV-G-CMV-blank) was used as a control (Abmgood, aavp002). In order to knockdown FGF-AS, lentivirus (LV) expressing shRNA targeting rat NUDT6 mRNA was injected into the hippocampus (Smart Vector Lentiviral Rat NUDT6 shRNA, Clone id: V3SVRN00_21213469, antisense sequence: AACCCTTCCCTGTGTCCG, Dharmacon). Non-targeting shRNA (with no complementarity to any known mammalian gene) expressing LV was used as control (Smart Vector Non-targeting hCMV-TurboGFP-Control, catalog# S-005000-01, Dharmacon).

### SURGICAL PROCEDURES

Rats were anesthetized with ketamine (50-100 mg/kg) and xylazine (5-10mg/kg) prior to surgical procedures. For icv injections, following the application of anesthesia 26 gauge guide cannulas (Plastics One Inc., VA. USA) were implanted into the left lateral ventricle (coordinates from bregma: anteroposterior −1.4, mediolateral +1.6, dorsoventral −2.5) and anchored to the skull with dental cement for. Five to seven days after surgery, a 33 gauge injector cannula was inserted extending 1 mm below the tip of the guides. The microinjection cannula was connected by PE-50 tubing to a Hamilton syringe mounted on a syringe pump (World Precision Instruments, FL. USA). Rats were microinjected with either pEGFP-N1 plasmid expressing full-length rat FGF-AS gene or in vivo JetPEI mix. Injections were performed at a rate of 1μl/min (total volume of injection=7 μl) for 14 days.

For viral injections, either NUDT6-AAV or NUDT6-shRNA-LV were injected bilaterally into the hippocampus (coordinates from bregma: anteroposterior −5.0, mediolateral +/-3.5, dorsoventral −4.0) by two micro-injectors (23 gauge), which were connected by PE-50 tubing to two Hamilton syringes mounted on a syringe pump (World Precision Instruments, FL. USA). Injections were performed at a rate of 0.25μl/min for 4 minutes; the microinjectors were left at the injection site for 5 minutes to allow diffusion of the virus then removed slowly in 5 minutes. The cranial hole was filled with bone wax and the skin was sutured with 6.0 silk.

### BEHAVIORAL TESTS

Behavioral tests were conducted 4 weeks after the injections of AAV or LV in order the constructs to express and exert their functions. pEGFP-N1 plasmid expressing full-length rat NUDT6 was injected for 14 days, and sucrose preference test and elevated plus maze test were conducted on the 11th and 13th day of injections, respectively; whereas forced swim test was evaluated 1 day after the last injection.

#### Sucrose Preference Test

The first day of the test, rats are habituated to consume sucrose by placing 2 bottles both filled with 1% sucrose solution. Then on the next day, one of the bottles is replaced with tap water, and 24 hours later total consumption of water and sucrose solutions is measured. For the repeated icv NUDT6-EP or vehicle-injected groups, animals were given access to 2 bottles, one filled with 1% sucrose and the other one filled with tap water for 2 hours after 12 hours of water deprivation. Total water and sucrose solution consumption were measured at the end of 2 hours. Sucrose preference is calculated as the percentage of sucrose consumption to total fluid consumption.

#### Elevated Plus Maze (EPM)

The rats are placed at the center of an elevated plus maze facing the closed arm. All four arms are 55 cm high, 35 cm long and 8 cm wide, with walls surrounding the closed arms that are 51 cm high. The frequency of entrance into the center, closed arms, open arms and distal half of open arms as well as time spent in each compartment are recorded for 5 minutes and analyzed with Ethovision XT v.8 software (Noldus Information Technology).

#### Forced Swim Test (FST)

The rats are placed in transparent cylinders (diameter=26 cm, height=43 cm), filled with water at 22-24°C for 15 minutes and 5 minutes on the first and second days of the experiment, respectively. During the second day, the time spent swimming and climbing and the immobility time are recorded and then manually scored with Ethovision XT v.8 software by an investigator blinded to groups.

### WESTERN BLOTTING

Following the euthanasia of the rats, the hippocampi were rapidly dissected on ice after. All samples were homogenized in RIPA buffer (NaCl, Triton-100-X, sodium deoxycholate, 1.5 M Tris Buffer [pH= 8.0], %10 SDS). Protein concentration was measured by BCA protein assay kit (Pierce) and equal amount from each sample was loaded and separated on 4-12 % Bis-Tris SDS gels (Novex, Invitrogen). The proteins were transferred to PVDF membranes (Novex, Invitrogen), which were blocked in 5% dry milk powder in TBS-T for 1 hour and then incubated in primary antibody solutions (Rabbit anti NUDT6, Proteintech 1:800; mouse anti FGF2, Millipore, 1:500; mouse anti ß-actin, Sigma, 1:7500; Rabbit anti ERK1/2, 1:1000, CST; Rabbit anti Akt 1:800, CST; Rabbit anti phospho-ERK1/2, 1:1000, CST; Rabbit anti phospho-Akt, 1:1000, CST; Rabbit anti S100A9, 1:200, CST). After treatment with the secondary antibody (anti-rabbit HRP-conjugated, 1:1000 or anti-mouse HRP-conjugated, 1:1000, Sigma), membranes were incubated in chemoluminescent (SuperSignal West Femto, Pierce) for 5 minutes and then imaged with Kodak 4000MM Image station. The band intensities were measured by Image J software (NIH). The optical intensities of all bands were normalized to that of beta-actin. To be able to compare band intensities on more than one membrane, each membrane was loaded with the same 2 samples, and the mean of the band intensities of these samples were used to normalize the intensities of the other bands on the same membrane.

### IMMUNOFLUORESCENT STAINING

For immunofluorescent staining, rats were first perfused with 4% paraformaldehyde (PFA) under deep anesthesia and their brains were collected. For each hippocampus three 40 μm thick sections that are 200 μm apart were chosen for immunolabeling. Sections were blocked in 10% NGS at room temperature (RT) and incubated in primary antibody (Anti S100A9, 1:200, CST; Anti Iba-1, 1:200, Wako; Anti NeuN, 1:200, Millipore; Anti NF-κB p65, 1:200, CST) at 4 °C overnight and in secondary antibody (Goat Anti-rabbit 1:200, Invitrogen) at RT for 1 hour. For double immunofluorescent stainings sections were blocked in 10% NGS at RT and incubated in primary antibodies simultaneously (Anti S100A9, 1:200, CST-Anti NF-κB p65, 1:200, SantaCruz; Anti NUDT6; ProteinTech, 1:200-Anti NF-κB p65, 1:200, SantaCruz; Anti NUDT6; ProteinTech, 1:200-Anti FGF2; Thermo Fisher, 1:200) at 4 °C 48 hours and in secondary antibodies simultaneously (Goat Anti-mouse 1:200, Invitrogen, Goat Anti-Rabbit 1:200, Jackson) at RT for 1 hour. There is an antigen retrieval step (80°C in citrate buffer) for all NF-κB double staining before blocking step. For NUDT6-S100A9 double staining; sections were blocked in %10 NGS at RT and incubated in anti NUDT6 antibody (1:200, ProteinTech) at 4 °C 48 hours and in secondary antibody (Goat Anti-Rabbit 1:200, Jackson) at RT for 1 hour. S100A9-NeuN or S100A9-Iba1 co-stainings were performed by incubating the sections in both primary antibodies which are labeled with fluorescent F(ab)_2_ fragments at 4 °C for 20 hours. Images from hippocampus were taken with 25X or 40X magnification with confocal microscope (Leica). An investigator blinded to the groups performed analyses of the labeled sections. Upon imaging, corrected total cell fluorescence (CTCF) was calculated for S100A9 labeling in the hilar and dentate gyrus neurons, respectively. Briefly, fluorescence intensity of S100A9 labeling was measured in NeuN positive neurons (ROI) via ImageJ from which background fluorescence was substracted. CTCF is calculated by the following formula: Fluorescence intensity (ROI)- (area of the selected cell x mean fluorescence of background readings). The nuclear translocation of NF-κB-p65 was calculated by the ratio of dentate gyrus cells positive for nuclear NF-κB staining to the total number of Hoechst positive dentate gyrus cells (around 200 cells per image). We counted and averaged NF-κB positive nuclei in 3 sections, 200 μm apart. Iba-1 positive microglia were imaged in Z-stacks and counted and averaged in maximum projection images obtained from three, 200 μm apart, sections. For S100A9 and Iba-1 co-labeling, ~25 μm Z-stacks (1024×1024 pixels, ~370×370 μm2) with isotropic voxel spacing were processed using FIJI/ImageJ 2.1.0/1.53f. for background subtraction (Rolling ball Radius of 50 pixels and 5 pixels for red and green channel, respectively).

#### Quantitative RT-PCR

Hippocampi were dissected and stored in RNAlater solution (QIAGEN) at −20 C until RNA isolation. Total RNA was extracted from the hippocampi with RNeasy mini kit (QIAGEN) as per manufacturer’s instructions. For cDNA synthesis 500 ng of total RNA was used. cDNAs were synthesized with random hexamer primers with RevertAid First Strand cDNA Synthesis Kit (Thermo Fisher Scientific, K1621) according to instructions and stored at −20°C. Quantitative RT-PCR was performed with Taqman probe-based technology. Taqman gene expression master mix (ABI, Foster city, CA, 4369016), FAM-TAMRA labeled Taqman probes for rat FGF-AS transcript (Assay ID: Rn00710461_m1), rat beta-actin gene (Assay ID: Rn01775763_g1), rat FGF2 gene (Taqman probe: CGT CCA TCT TCC TTC ATA GCC AG; forward primer: GGA GTT GTG TCC ATC AAG; reverse primer: CAC TCT TCT GTA ACA CAC TTA) and rat S100A9 gene gene (Taqman probe:TGC TGC GCT CCA GCT GAG ATC C; forward primer: GCT TTG AGC AAG AAG ATG GCT; reverse primer: CCT TGT TCA GGG TGT TCA GGA) were used. PCR was carried out in triplicates in ABI OneStep Q RT-PCR machine (ABI). Thermal cyclic conditions were as follows: 50°C for 2 min, 95°C for 10 min followed by 40 cycles of 95°C 15 s, 60°C for 1 min. The relative expression values were calculated with ΔΔCt method.

### PROTEOMIC ANALYSIS

Hippocampi that were dissected 8 weeks after the injection of either NUDT6 overexpressing AAV (n=5) or Blank AAV (n=6) were sent to Acibadem University Department of Biochemistry for proteomic analysis by using Liquid Chromatography Tandem Mass Spectrometry (LC-MS/MS). The raw data obtained from LC-MS/MS was processed and compared between groups. Principal Component Analysis was performed and proteins with significant expression differences were determined. The network and pathway analysis were performed using online bioinformatics websites (Panther, String). The activated or inhibited proteins were determined by their relative activation z-scores.

### STATISTICAL ANALYSES

Sample sizes were selected based on similar studies in the literature; no statistical analysis was conducted to predetermine sample size. Data was analyzed by SPSS 22. Kolmogorov Smirnov Test was used to test for the normality of the distribution. Variables that were normally distributed were compared with student t-test, whereas those that were not normally distributed were compared with Mann Whitney-U test. All statistical tests were conducted two-sided. Data are presented as mean ± S.E.M.

## ACKNOWLEDGEMENTS

This project was supported by the Scientific and Technological Research Council of Turkey, grant number: SBAG 110S481; L’Oréal-UNESCO For Women in Science Award to EEK; Hacettepe University Research Projects Management System, grant number: THD-2016-11687, and Science Academy Young Scientists Award Program to EEK. We thank Dr. Paul Murphy (Dalhousie University, Canada) for his generosity in providing the rat NUDT6 expressing plasmid, and Mesut Firat and Sinem Yilmaz-Özcan for their expert help in technical issues. We thank Dr Cortney Turner and Aiden Houcek for their valuable opinions on the manuscript. Authors declare no competing financial interests. Graphical depictions were prepared at Biorender.com.

## AUTHOR CONTRIBUTIONS

Design and conception of the study: EEK; acquisition and analysis of data: EEK, BU, FOH, MY, ECE, ABV and KB; scientific discussions and interpretation of data: BU, EEK, TD, YA, KB; drafting the manuscript: BU, EEK and TD

