## Supplemental Data for "NUDT6, the Antisense Protein of FGF2 Gene, Plays a Depressogenic Role by Promoting Inflammation and Suppressing Neurogenesis without Altering FGF2 Signaling"

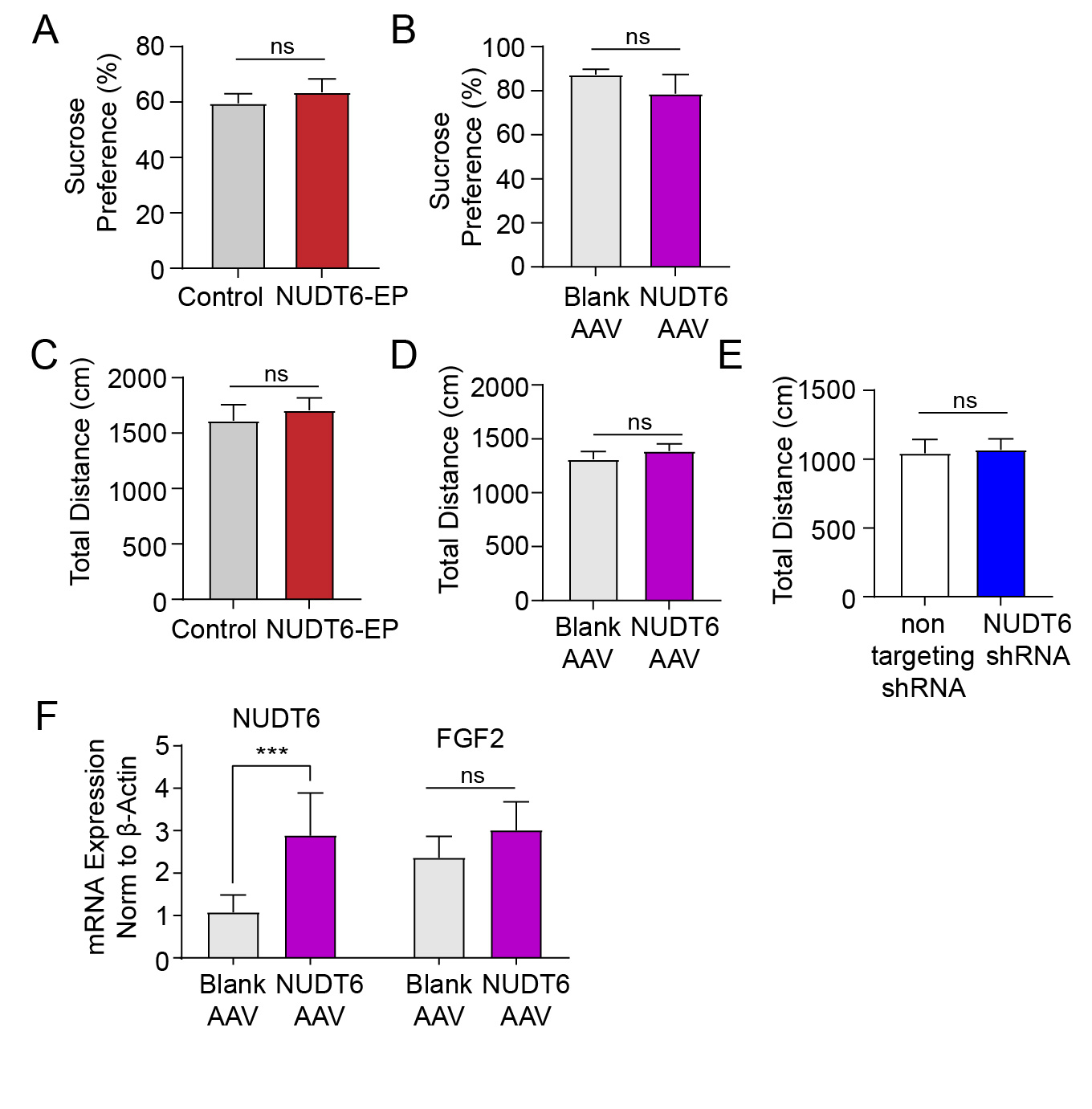

**Supplementary Figure 1. A-B**. NUDT6 overexpression by neither icv injection of NUDT6-EP nor intra-hippocampal injection of NUDT6-AAV changed sucrose preference when compared to their respective control groups **C-D.** Neither repeated icv NUDT6-EP injections nor intrahippocampal NUDT6-AAV injections changed the total distance moved in EPM when compared to their relevant control groups. **E.** Intrahippocampal NUDT6-shRNA injections did not change total distance moved in EPM when compared to non-targeting shRNA injected group. **F.** Hippocampal NUDT6 overexpression significantly increased NUDT6 mRNA levels but did not change FGF2 mRNA levels normalized to β-actin.

FGF2: fibroblast growth factor 2EPM: elevated plus maze, NUDT6-AAV: NUDT6 expressing adeno-associated virus, NUDT6-EP: NUDT6 expressing plasmid, NUDT6-shRNA: NUDT6 short hairpin RNA, icv: intracerebroventricular. Significance levels were stated as follows: ***p < 0.001. ns denotes non-significance.

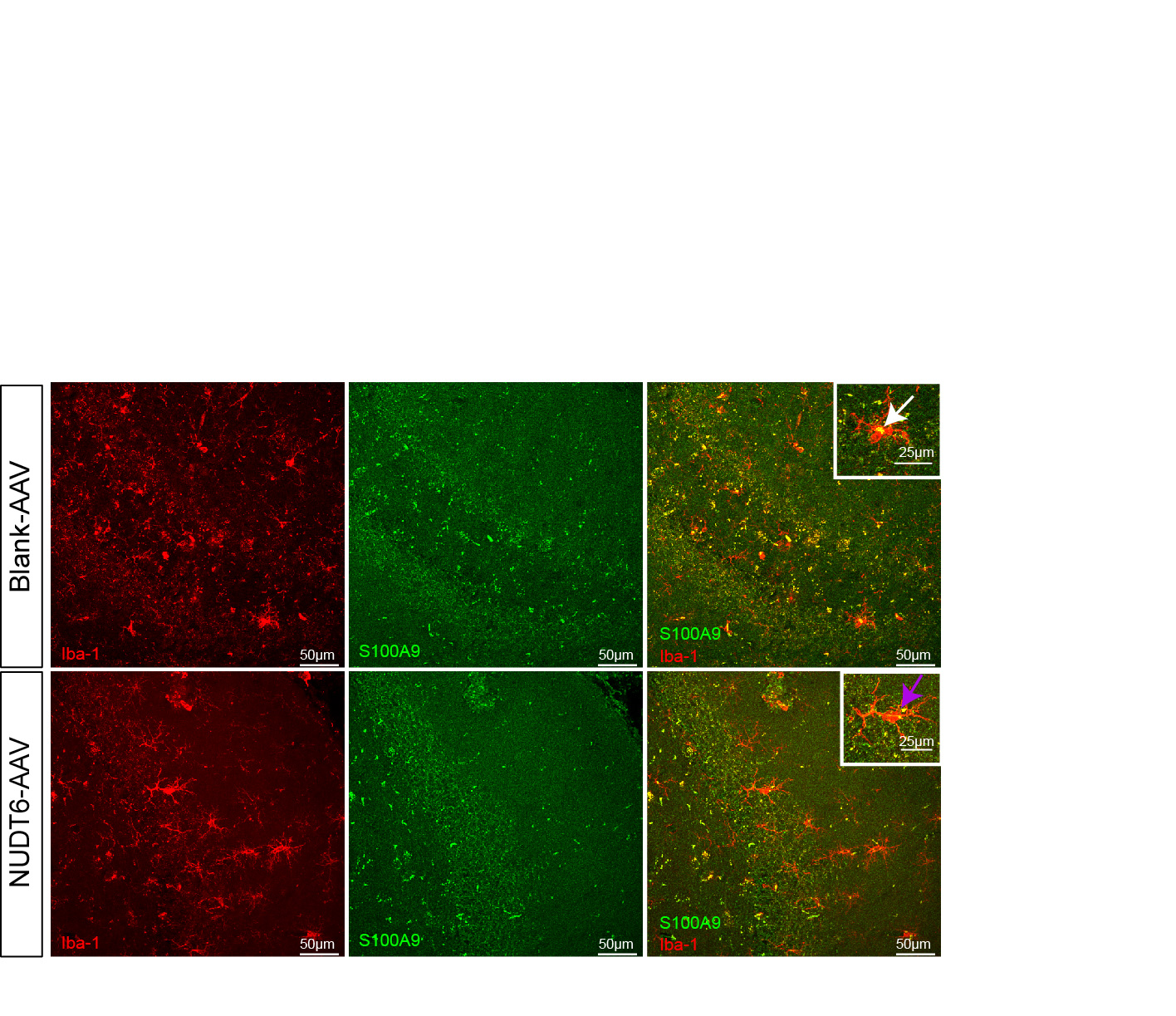

**Supplementary Figure 2.** Colocalization of S100A9 with a microglial cell marker, Iba1 showed that S100A9 was expressed in all microglial cells labelled with Iba1. Microglia were labeled with Iba1 (red) and S100A9 was labeled with green. Figure inset show a representative microglial cell expressing S100A9. Scale bar: 50 μm and 25 μm for the inset

|  | **PROTEIN** | **FOLD CHANGE** |  |
| --- | --- | --- | --- |
| 1 | Chloride intracellular channel protein 6 (CLIC6_RAT) | 46.4 | increase |
| 2 | Protein S100-A9 (S10A9_RAT ) | 32.1 | increase |
| 3 | Alpha-1-antiproteinase (A1AT_RAT) | 4.2 | increase |
| 4 | Transthyretin (TTHY_RAT) | 3.9 | increase |
| 5 | Ig gamma-2A chain C region (IGG2A_RAT) | 3.3 | increase |
| 6 | Fibrinogen gamma chain (FIBG_RAT) | 2.8 | increase |
| 7 | Alpha-1-macroglobulin (A1M_RAT) | 2.6 | increase |
| 8 | Band 3 anion transport protein (B3AT_RAT) | 2.6 | increase |
| 9 | Erlin-2 (ERLN2_RAT) | 2.3 | increase |
| 10 | Nucleobindin-1 (NUCB1_RAT) | 1.9 | increase |
| 11 | Tropomyosin beta chain (TPM2_RAT) | 1.9 | increase |
| 12 | Glutathione peroxidase 1(GPX1_RAT) | 1.8 | increase |
| 13 | Cancer-associated gene 1 protein homolog (CAGE1_RAT) | 1.8 | increase |
| 14 | Hemoglobin subunit beta-1 (HBB1_RAT) | 1.8 | increase |
| 15 | Carbonic anhydrase 1 (CAH1_RAT) | 1.8 | increase |
| 16 | Serotransferrin (TRFE_RAT) | 1.7 | increase |
| 17 | Hemoglobin subunit beta-2 (HBB2_RAT) | 1.6 | increase |
| 18 | TSC22 domain family protein 1 (T22D1_RAT) | 1.6 | increase |
| 19 | Hemoglobin subunit alpha-1/2 (HBA_RAT) | 1.6 | increase |
| 20 | Vimentin (VIME_RAT) | 1.6 | increase |
| 21 | Fibrinogen alpha chain (FIBA_RAT) | 1.5 | increase |
| 22 | Keratin_ type I cytoskeletal 18 (K1C18_RAT) | 1.5 | increase |
| 23 | *Vesicle-associated membrane protein 1 (VAMP1_RAT)* | *1.5* | *decrease* |
| 24 | Vinculin (VINC_RAT) | 1.4 | increase |
| 25 | Stress-induced-phosphoprotein 1( STIP1_RAT) | 1.4 | increase |
| 26 | *Vesicular inhibitory amino acid transporter (VIAAT_RAT)* | *1.4* | *decrease* |
| 27 | Mitochondrial fission 1 protein (FIS1_RAT) | 1.4 | increase |
| 28 | *Sodium/potassium-transporting ATPase subunit beta-2 (AT1B2_RAT)* | *1.3* | *decrease* |
| 29 | Core histone macro-H2A.1 (H2AY_RAT) | 1.3 | increase |
| 30 | Solute carrier family 12 member 5 (S12A5_RAT ) | 1.3 | increase |
| 31 | 6-phosphogluconolactonase (6PGL_RAT ) | 1.3 | increase |
| 32 | Histone H1t (H1T_RAT) | 1.3 | increase |
| 33 | NADH dehydrogenase [ubiquinone] iron-sulfur protein 6_ mitochondrial (NDUS6_RAT) | 1.3 | increase |
| 34 | Nucleoprotein TPR (TPR_RAT) | 1.3 | increase |

**Supplementary Table 1.** Proteomic comparison of NUDT6-AAV and blank-AAV injected hippocampal samples showed a significant increase in the expression of 31 proteins whereas a decreased expression of 3 proteins**.**
